# StrucTTY: An Interactive, Terminal-Native Protein Structure Viewer

**DOI:** 10.64898/2026.03.17.712308

**Authors:** Luna Sung-eun Jang, Sooyoung Cha, Martin Steinegger

## Abstract

Terminal-based workflows are central to large-scale structural biology, particularly in high-performance computing (HPC) environments and SSH sessions. Yet no existing tool enables real-time, interactive visualization of protein backbone structures directly within a text-only terminal. To address this gap, we present StrucTTY, a fully interactive, terminal-native protein structure viewer. StrucTTY is a single self-contained executable that loads mulitple PDB and mmCIF files, normalizes three-dimensional coordinates, and renders protein structures as ASCII graphics. Users can rotate, translate, and zoom in on structures, adjust visualization modes, inspect chain-level features and view secondary structure assignments. The tool supports simultaneous visualization of up to nine protein structures and can directly display structural alignments using Foldseek’s output, enabling rapid comparative analysis in headless environments. The source code is available at https://github.com/steineggerlab/StrucTTY.

Key Messages

- Real-time, interactive protein structure visualization directly within text-only terminals
- ASCII-based, depth-aware rendering of PDB and mmCIF backbone structures
- Multi-structure comparison with direct application of Foldseek alignment transformations
- Designed for headless workflows on remote servers and HPC systems

## Introduction

Visualization remains fundamental to structural biology. Even with advances in sequence analysis and structure prediction, researchers continue to rely on direct structural inspection to interpret folding patterns, domain organization, and structural similarity. Consequently, visualization platforms such as PyMOL, UCSF ChimeraX, VMD, and Mol* (1; 2; 3; 4) are widely used in structure-based research workflows.

However, modern structural bioinformatics increasingly operates in high-performance computing (HPC) environments, remote servers, and text-only SSH sessions. In these contexts, graphical interfaces are often unavailable or impractical. Researchers must either transfer files to local machines for inspection or interpret raw coordinate files directly, both inefficient when handling large numbers of structures. Despite the centrality of terminal-based workflows in computational biology, no existing tool enables real-time, interactive visualization of protein structures directly within a text-only terminal. While limited ASCII rendering attempts exist, they lack essential capabilities required for structural analysis, including consistent 3D normalization, depth-aware projection, multi-chain visualization, and multi-structure comparison.

To address this gap, we developed StrucTTY, a fully interactive, terminal-native protein structure viewer. StrucTTY is distributed as a single self-contained executable capable of loading multiple PDB and mmCIF files, normalizing three-dimensional coordinates, and rendering backbone structures as depth-aware ASCII graphics in real time.

StrucTTY is implemented for POSIX-compatible terminal environments and runs natively on Linux systems. Because it depends only on standard terminal capabilities and does not require a graphical subsystem, it can be executed on most Unix-like platforms, including macOS terminals and Windows environments providing POSIX-compatible shells such as Git Bash or MinTTY. In addition, Linux terminal environments available on mobile devices (e.g., via terminal emulation applications) allow StrucTTY to run on smartphones and tablets. This broad platform compatibility reinforces StrucTTY’s portability across desktops, remote servers, cloud systems, and lightweight devices.

StrucTTY is designed specifically for headless environments and HPC workflows. It supports interactive navigation (rotation, translation, zooming), multi-structure comparison, secondary structure highlighting, and direct application of alignment transformations generated by Foldseek (5). By integrating structural visualization directly into terminal workflows, StrucTTY enables rapid exploratory analysis without graphical dependencies.

## Interface

Figure 1 illustrates the StrucTTY interface, which consists of two primary components: a rendering screen and an information panel.

**Figure 1.**
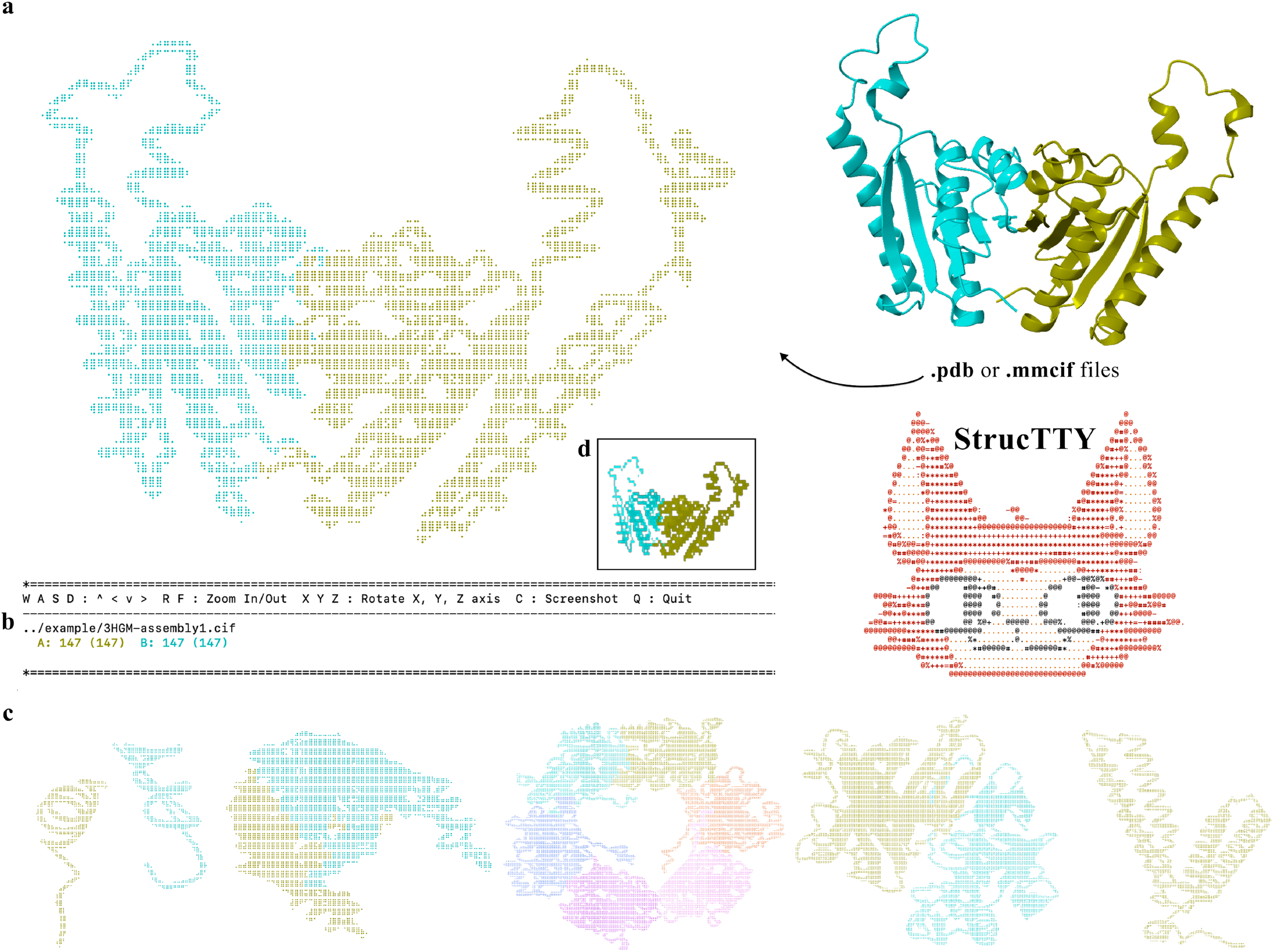
Protein backbone structure visualization with StrucTTY. (a) 3HGM bioassembly mmCIF file rendered by StrucTTY. (b) The interactive command and chain information panel displayed alongside the structure view. Panels (a) and (b) together represent the actual output shown in a user’s terminal during interactive use. (c) Representative examples of protein structures visualized with StrucTTY, shown from left to right: 5W96, 3HGM, 9N47, 3OAG, and A0A233SAX3. (d) A rasterized image generated by the -c screenshot command, which saves the current terminal visualization as a PNG file. This panel is not part of the live terminal display but represents exported output for downstream use.

The rendering screen displays C_*α*_ backbone coordinates projected from three-dimensional space into a depth-aware ASCII representation. The structure viewer and panel sizes dynamically adapt to the terminal window dimensions. The visualization dynamically reflects the current viewing parameters, including orientation, zoom level, rendering style, and selected chains.

The information panel provides structural metadata and usage guidance, including chain identifiers, residue counts, and available commands. This dual-layout design allows users to inspect structural features while maintaining contextual information within a single terminal view.

### Basic Controls

StrucTTY supports real-time manipulation of structures through keyboard input:

- **Translation**: Move the structure using `W/A/S/D`.
- **Rotation**: Rotate around the X, Y, and Z axes using `X, Y`, and `Z`.
- **Zoom**: Adjust scale using `R` (zoom in) and `F` (zoom out).
- **Screenshot**: Export the current frame as PNG or text using C.
- **Protein Summary Panel**: Display chain identifiers and residue counts.

### Visualization Options

StrucTTY provides configurable visualization options to enhance interpretability across diverse terminal environments (Figure 2).

**Figure 2.**
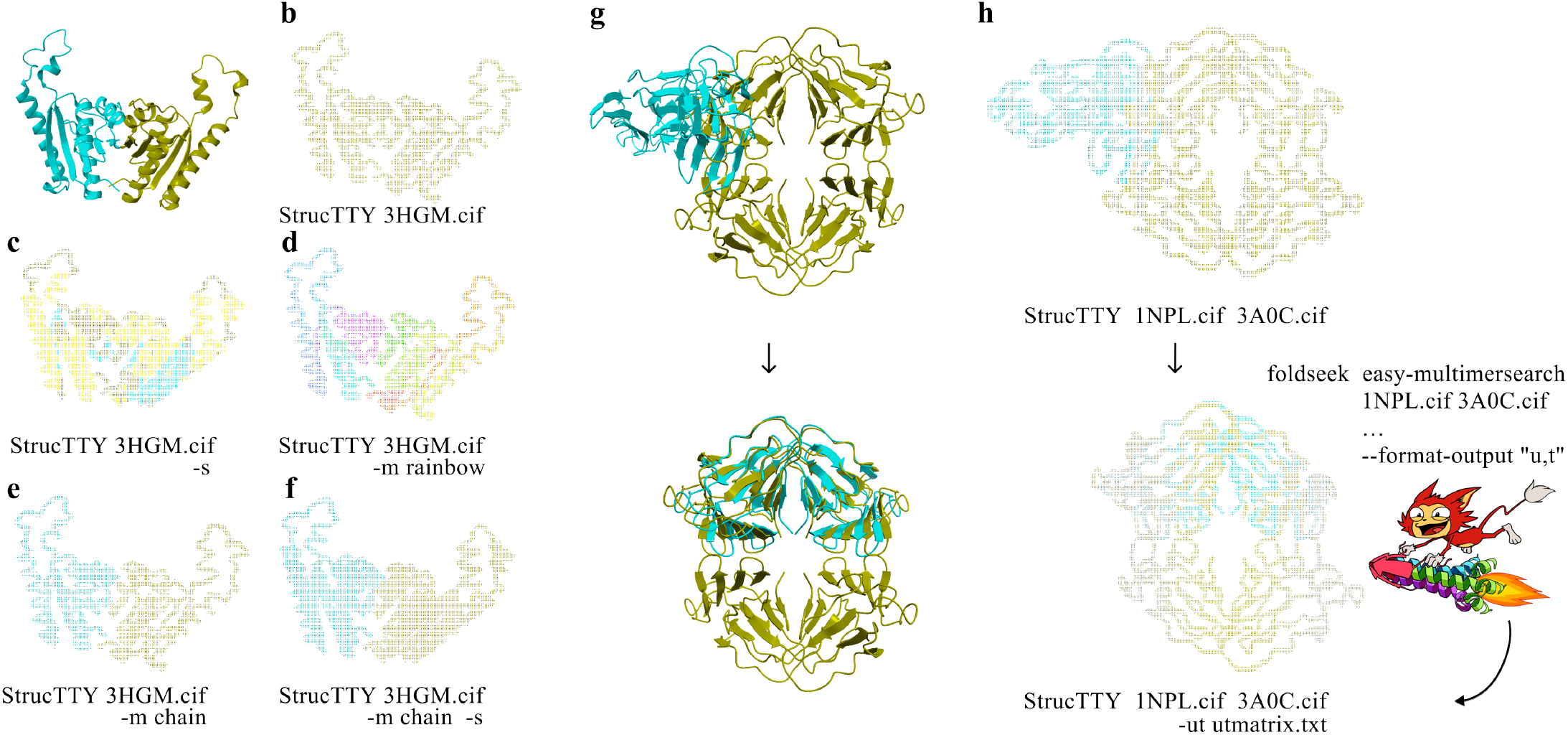
Visualization and alignment-based structural comparison in StrucTTY (a) Conventional 3D ribbon representation of 3HGM bioassembly cif file using ChimeraX shown for reference. (b-f) StrucTTY outputs of 3HGM bioassembly cif file with different paramters for coloring and visualizing secondary structures. (g) Conventional ribbon representations of 1NPL and 3A0C bioassembly mmCIF files using ChimeraX shown for the references(above) and rotated and translated result based on foldseek alignment(below). (h) StrucTTY result with the default parameter(above) and with foldseek alignment result as an input utfile from Foldseek(below).

- **Color Modes**: Supports `-m rainbow, -m chain, and -m default` to distinguish residues or chains.
- **Secondary Structure Display**: Highlights helices and sheets with shape and color using `-s`. When annotations are absent, secondary structures can be inferred from backbone geometry.
- **Chain Selection**: Selectively render chains via a mapping file passed to `-c`.

### Expert Options

Advanced functionality supports multi-model comparison and integration with structural alignment tools (Figure 2g,h).

- **Multiple Proteins**: Up to nine structures can be loaded simultaneously and toggled using keys `0--9`.
- **Foldseek Transformation Matrices**: Alignment transformations can be applied using the `-ut` option. The input file must contain the protein index, nine rotation values, and three translation values.

Overall, the StrucTTY interface is designed to balance simplicity and expressiveness within a purely text-based environment. By combining real-time interaction, flexible visualization options, and integrated structural context, it enables efficient inspection and comparison of protein structures without disrupting terminal-based workflows.

## Use Case and Benchmarking

Established visualization platforms including PyMOL, UCSF ChimeraX, VMD, and Mol* provide high-quality graphical rendering, advanced analysis features, and extensive scripting capabilities. PyMOL excels in publication-quality rendering, ChimeraX in GPU-accelerated visualization and cryo-EM analysis, VMD in large-scale molecular dynamics, and Mol* in browser-based accessibility.

However, these tools depend on graphical environments or web browsers. In headless HPC systems or remote SSH sessions, they require additional configuration or local file transfer for visualization. StrucTTY addresses a complementary use case: lightweight, interactive structural inspection directly within text-only terminals. Rather than replacing full-featured graphical viewers, StrucTTY extends structural visualization into computational environments where graphical tools are impractical.

To evaluate performance, we benchmarked StrucTTY across representative protein structures of varying sizes (Table 1). Benchmarks were executed on a SLURM-managed Linux compute node using a single CPU core and 4 GB of memory (`srun -c 1 --mem=4G`). Each structure was evaluated in benchmark mode five times under identical conditions (`--show_structure, --no_panel`). Performance metrics—including load time, time-to-first-frame (TTFF), per-frame render time, and input-to-frame latency—were recorded using `std::chrono::steady_clock`. Mean and 95th percentile values were reported across independent runs. Frame and latency statistics were obtained using a deterministic interaction protocol executed in benchmark mode. Each run consisted of 200 warm-up iterations followed by 2000 timed iterations to minimize initialization and caching effects.

**Table 1.**
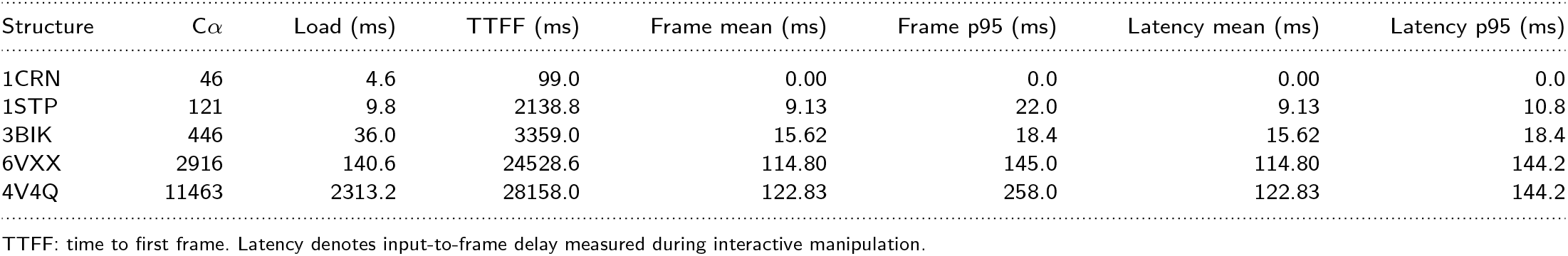
Benchmark performance of StrucTTY across representative protein structures of varying sizes. C*α* count corresponds to the number of rendered residues. Values represent the mean across 5 independent runs; p95 denotes the 95th percentile.

| Structure | Cα | Load (ms) | TTFF (ms) | Frame mean (ms) | Frame p95 (ms) | Latency mean (ms) | Latency p95 (ms) |
| --- | --- | --- | --- | --- | --- | --- | --- |
| 1CRN | 46 | 4.6 | 99.0 | 0.00 | 0.0 | 0.00 | 0.0 |
| 1STP | 121 | 9.8 | 2138.8 | 9.13 | 22.0 | 9.13 | 10.8 |
| 3BIK | 446 | 36.0 | 3359.0 | 15.62 | 18.4 | 15.62 | 18.4 |
| 6VXX | 2916 | 140.6 | 24528.6 | 114.80 | 145.0 | 114.80 | 144.2 |
| 4V4Q | 11463 | 2313.2 | 28158.0 | 122.83 | 258.0 | 122.83 | 144.2 |

TTFF: time to first frame. Latency denotes input-to-frame delay measured during interactive manipulation.

## Future works

Future development will focus on enhanced scalability and deeper integration with structural bioinformatics workflows. GPU-accelerated projection may improve performance for large assemblies. Closer coupling with structural search frameworks such as Foldseek could further streamline terminal-based workflows by directly connecting search, alignment, and visualization.

## Conclusion

StrucTTY introduces a terminal-native paradigm for interactive protein structure visualization. By eliminating graphical dependencies while preserving essential structural information, it enables direct structural inspection in headless computing environments.

Designed for portability and responsiveness, StrucTTY integrates naturally into large-scale computational pipelines and supports rapid exploratory analysis. Multi-structure visualization and alignment transformation support further extend its utility for comparative structural studies. Because it operates in POSIX-compatible terminal environments, StrucTTY runs natively on Linux systems and remains portable across macOS, Windows terminal shells, and lightweight terminal environments on mobile devices.

Together, these features establish terminal-native visualization as a practical and complementary approach within modern structural bioinformatics workflows.

## Acknowledgments

This work was supported by grants from the National Research Foundation of Korea [2020M3A9G7103933, RS-2021-NR061659, RS-2021-NR056571, RS-2024-00396026, RS-2024-00438101 to M.S.]; Samsung DS Research Fund; Creative-Pioneering Researchers Program; and Novo Nordisk Foundation [NNF24SA0092560].

